# Charge-engineered nanohybrid coatings for microbial fouling resistance

**DOI:** 10.64898/2026.11.18.767735

**Authors:** Ramesh Devadig, Venkata A.S. Kandadai, Kalimuthu Jawaharraj, Bharat K. Jasthi, Etienne Gnimpieba, Caine Shagla, Alan Dalton, James McHugh, Pulickel M. Ajayan, Manoj Tripathi, Venkataramana Gadhamshetty

## Abstract

Microbial attachment is the gateway to biofouling and microbially influenced corrosion. Yet biocides and passive barrier coatings do not actively regulate the charged metal– microbe interface at which adhesion begins. Here we report a charge-engineered coating— a graphene thin-film nanohybrid (gTFN) on nickel (Ni) forming an asymmetric metal– graphene–metal junction—that suppresses cell attachment through interfacial charge redistribution rather than soluble toxicity. Against the model sulfate-reducing bacterium Oleidesulfovibrio alaskensis G20, gTFN combines an impermeable graphene scaffold, an immobilized low-leaching Ag cap and a serrated topology, reducing microbial surface coverage by 98%, increasing charge-transfer resistance 47-fold and lowering corrosion current 167-fold relative to Ni. Surface-potential mapping and density functional theory reveal interfacial dipoles that pattern the surface potential, while imaging shows graphene wrinkles channeling residual attachment. Transcriptomic profiling reveals downregulation of biofilm-associated and energy-metabolism genes. These results establish interfacial charge redistribution as an active antifouling and anticorrosion mechanism for controlling microbial biointerfaces.

---

Microbial adhesion serves as the initial step toward biofouling.^1, 2^ and microbially influenced corrosion (MIC),^3, 4^ two interfacial processes that accelerate infrastructure degradation, metal leaching, and greenhouse gas emissions, contributing to multi-trillion-dollar global losses.^5^ Conventional protective coatings, including chromium-based paints, can slow corrosion but do not disrupt the physicochemical interactions that govern bacterial attachment. Nanostructured two-dimensional (2D) coatings offer barrier properties against biotic^6^ and abiotic corrosion.^3, 7^ However, they lack inherent control over microbial colonization. Surface-modification strategies, from hydrophilic hydration layers to hydrophobic interfaces, suppress adhesion by weakening polar and hydrogen-bondinginteractions.^8^ Yet, they do not actively regulate microbial–surface electrostatics. Antimicrobial nanoparticles are widely incorporated into antifouling coatings,^9^ but weak adhesion can cause uncontrolled release, raising concerns over antimicrobial resistance^10^ and ecosystem impacts.^11^ Their agglomeration, poor dispersion, and limited stability issues^12, 13^ further motivate immobilized, low-leaching architectures.

Localized electronic charge concentration and its associated electrostatic potential are increasingly recognized as regulators of protein adsorption, cell-surface interactions, and biofilm nucleation.^14, 15^ This suggests an opportunity to design coatings that modulate interfacial charge while presenting nanoscale and microscale textured topographies aligned with cell-attachment theory.^16^ For long-term performance, such coatings should suppress cell adhesion, protein secretion, and biofilm formation, particularly by corrosion-associated sulfate-reducing bacteria (SRB).^4^ Achieving this requires noninvasive materials that influence microbial behavior across phenotypic and genotypic levels without relying on soluble toxicity. A direct, solid-state route to such control is the metal–graphene–metal (MGM) junction: when graphene layers are sandwiched between two dissimilar metals, their distinct intrinsic work functions (eV) drive interfacial charge transfer, generating interface dipole layers and a built-in surface potential set by the choice of metals^17^ rather than by soluble chemistry. Here, we establish surface-charge topology as an active determinant of microbial colonization, distinct from prior graphene-based coatings that mitigate microbial attack largely through passive barrier effects^3, 7^.

We therefore present a graphene thin-film nanohybrid (gTFN) coating that realizes an asymmetric metal–graphene–metal (MGM) junction (Ni–graphene–Ag) to regulate microbial colonization through interfacial charge redistribution. gTFN is epitaxially assembled bottom-up: multilayer graphene (∼100 nm) is grown on polycrystalline Ni by chemical vapor deposition, with a buffer layer of a mixed-phase nickel carbide (Ni_3_C) between Ni and graphene layers^17, 18^. In the subsequent step, it is capped with an immobilized, low-leaching ∼3 nm Ag layer by pulsed laser deposition, forming a uniform and stable architecture. Multimodal analyses spanning electrochemistry, surface physics, machine-learning-assisted imaging, transcriptomic and first-principles calculations reveal strong antifouling and corrosion resistance, including a 98% reduction in bacterial attachment, a 47-fold increase in charge-transfer resistance and a 167-fold decrease in corrosion current. Transcriptomic profiling further reveals suppression of biofilm-associated and energy-metabolism genes. These responses support a layered defense mechanism in which interfacial charge redistribution, mapped by KPFM and DFT, is distinct from charge transfer resistance (R_ct_) measured by EIS. Across the Ni–graphene–Ag interface, this redistribution creates a surface-potential landscape that limits initial microbial contact, while the immobilized Ag cap and serrated graphene topology act on residual cells that reach the surface. Together, these findings establish surface-charge topology as an active design principle for low-leaching, charge-engineered coatings that suppress microbial colonization and downstream corrosion beyond passive barrier protection.

## Nanohybrid architecture, interfacial charge redistribution and gene expression

We first engineered a Ni–graphene–Ag architecture to test whether interfacial charge redistribution can regulate microbial attachment. gTFN was synthesized by chemical vapor deposition (CVD) of epitaxial multilayer graphene up to ∼100 nm thick on polycrystalline Ni (grown in place by carbon dissolution into Ni and precipitation on cooling—the solubility route, unlike self-limiting growth on Cu—rather than transferred), followed by pulsed laser deposition (PLD) of a ∼3 nm Ag cap (**Fig. 1a; see Material synthesis**). This stacked metal–graphene–metal (MGM) interface couples three electronically distinct layers: Ni, graphene and Ag. The junction is structurally asymmetric: graphene is covalently bonded to Ni through the Ni C buffer via Ni 3d–π interaction^19^, a stronger bond than the C–Ag contact beneath the Ag cap. Independently of this bonding asymmetry, the distinct intrinsic work functions of Ni, graphene and Ag drive interfacial charge transfer on contact, which together with the covalent Ni C–graphene redistribution sets the interface dipoles. Because the ∼100 nm graphene interlayer makes these interface dipoles act largely independently, local graphene topology—rather than a single through-stack field—patterns the surface potential that cells encounter (**Fig. 1c, d**).

**Fig. 1.**
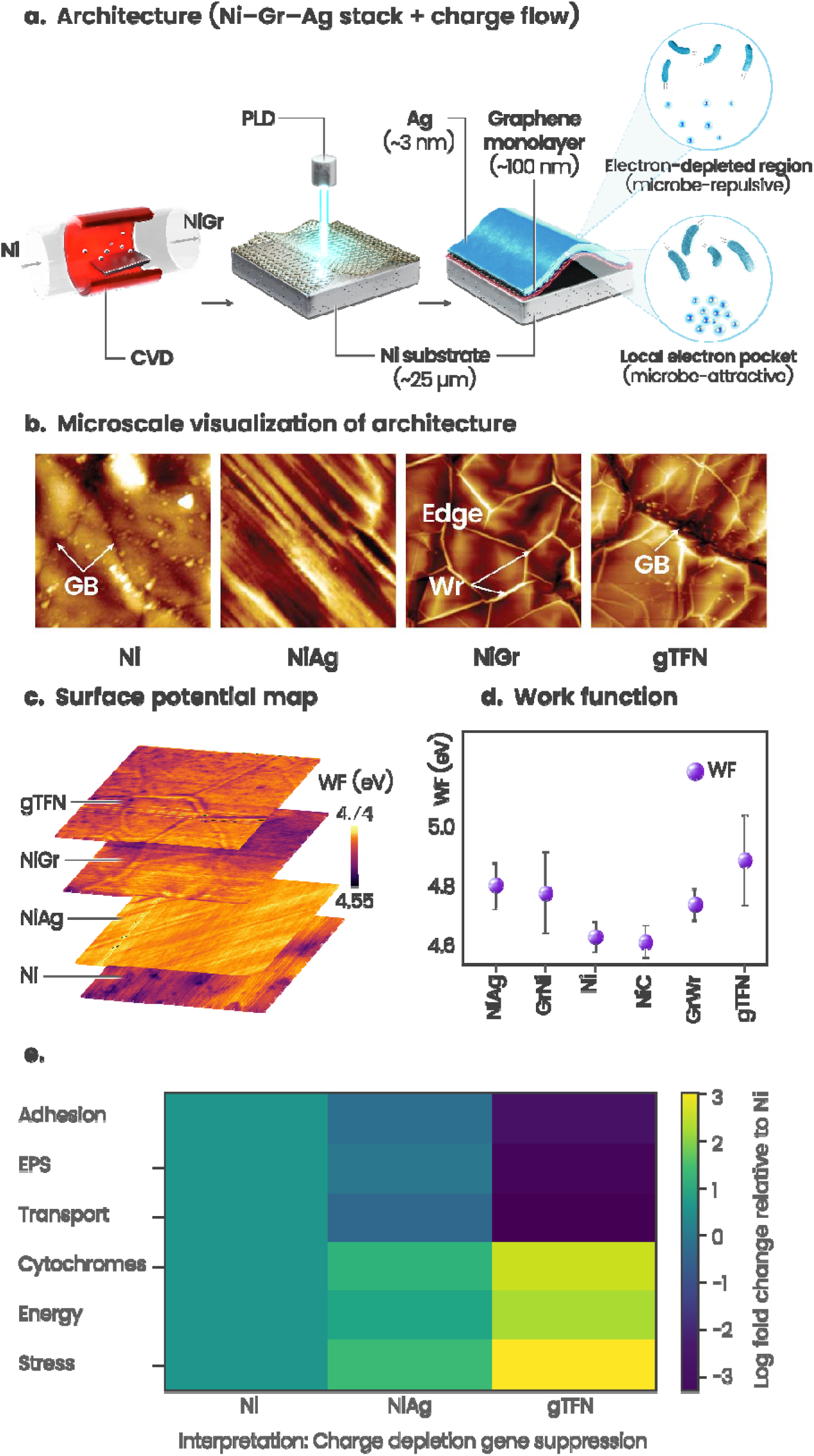
| Engineering gTFN and the interfacial charge landscape governing antifouling. (a) Schematic of gTFN synthesis by CVD-based deposition of graphene layers followed by PLD deposition of a nanosilver layer. (b) Atomic force microscopy (AFM) topography of bare Ni, NiAg, NiGr and gTFN surfaces, highlighting key structural differences: Ni exhibits exposed grain boundaries (GBs), while PLD-deposited Ag (NiAg) partially masks these defects. The epitaxial growth of graphene layers (NiGr) introduces a graphitic disordered structure of wrinkles and edge regions. The capping of Ag nanolayer in gTFN up to 3 nm thickness conforms over flat and disordered regions. (c) Surface potential map revealing the alteration of work function (eV) after addition of the graphitic layer and silver coatings, respectively. (d) Average WF values across different substrates and graphitic disordered structures influence microbial interactions. (e) Heat map of Ni, NiAg and gTFN showing surface-induced transcriptomic reprogramming.

Sandwiched between Ni and Ag, graphene acts as an intermediate electron reservoir and is bonded to Ni through a nickel carbide buffer layer that is less dense than the graphene layers^17^. The work-function mismatch across these layers promotes charge redistribution through multiple interface dipole layers (IDL), supporting charge-carrier-driven surface reactions,^20^ that are expected to influence microbial attachment and biofilm formation.

The architecture also creates a spatially heterogeneous surface for microbial contact. Graphene wrinkles and edges introduce localized electron-rich regions, whereas the basal plane remains comparatively charge-depleted (**Fig. 1a)**. This contrast provides a basis for channeling residual microbial attachment to wrinkle domains while limiting attachment across the broader basal-plane surface. The ultrathin Ag cap conforms to this graphene morphology while providing an immobilized, low-leaching antimicrobial component (see Supplementary Methods **S1** and Supplementary **Fig. S1**). ^21^ It may also help mask topographic defects in Ni and graphene-covered Ni, including grain boundaries, pit holes and the native Ni oxide layer (**Fig. 1b).** Consistent with this design, surface-potential mapping across Ni, NiAg, NiGr and gTFN shows a progressive change in the surface charge landscape (**Fig. 1c**), and work-function analysis indicates that graphene and the Ag cap increase the work function relative to Ni (**Fig. 1d**), a condition expected to disfavor microbial attachment initiation.^22, 23, 24^

This charge-led interpretation is consistent with prior reports showing that metal–graphene interfaces with dissimilar electronegativities can catalyze reactions or facilitate antimicrobial activity. Graphene on metallic (Cu) and semiconductive (Ge) substrates exhibits greater antimicrobial activity than on insulating substrates, owing to dipole formation and interfacial charge redistribution.^25, 26^

In the present architecture, Ag ions in the cap can accept electrons from microbial cell membranes and disrupt physiological activity,^27^ while extracted charge is redistributed across the Ag–graphene–Ni interfaces through multidipole layers^28^. Thus, gTFN integrates graphene structural stability and conductivity with an immobilized Ag cap, producing a low-leaching interface that combines charge redistribution, surface topology and antimicrobial activity.

We next examined whether this charge-engineered interface altered the OA-G20 transcriptional response. Transcriptomic profiling was performed using the corrosion-associated SRB OA-G20.^29^ We investigated interfacial bacterial responses across substrates, focusing on processes associated with protein folding and cell-wall biosynthesis. A total of 371 and 305 differentially expressed genes were identified in OA-G20 exposed to gTFN and NiAg, respectively, relative to Ni, as compiled in the heat map in **Fig. 1e**, based on **Supplementary Tables S1a, S1b, and S1c**. The heat map indicates reduced expression on gTFN across adhesion-related, transporter, flagellar/motility, exopolysaccharide (EPS)/biofilm, stress-adaptation and polyamine-transport categories. Because NiGr isolates the graphene-induced charge contribution in the absence of Ag, the NiGr, NiAg and gTFN series distinguishes charge-governed inhibition from Ag-mediated effects and establishes NiGr as the silver-free control for the charge mechanism.

The gene-level response supports impaired attachment on gTFN. Adhesion-related and transporter genes remained active on NiAg, consistent with partial microbial colonization (Supplementary **Fig. S2a**, NiAg vs Ni). In contrast, gTFN exposure significantly downregulated the spermidine/putrescine transporter PotA (Dde_3670), peptidoglycan-associated proteins (Dde_3633, Dde_3602) and flagellar/motility components (Dde_1654, Dde_3151). These genes contribute to biofilm maturation, EPS stabilization and motility-driven adhesion. Their suppression suggests that OA-G20 cells are less able to establish stable attachment on gTFN.

At the same time, gTFN activated a stress-adaptation response rather than a colonization program. Upregulated genes included stress-related enzymes, such as D-alanyl-D-alanine dipeptidase (Dde_1551), and redox proteins, including hydrogenase, consistent with metabolic stress (**Supplementary Fig. S2b**, gTFN vs Ni). Direct comparison with NiAg showed that gTFN induced stronger metabolic reprogramming, suppressing structural and transporter genes while activating stress-adaptation pathways (Supplementary **Fig. S2c**, gTFN vs NiAg).

The Ag–graphene multilayer combination in gTFN yielded 374 DEGs relative to NiAg, indicating additional gene-regulatory effects. Compared with Ni, the spermidine/putrescine import ATP-binding protein gene potA (Dde_3670), which supports polyamine uptake, cell proliferation and ion homeostasis, was downregulated 3.3-fold in gTFN and 2.84-fold vs. NiAg. Thus, gene ontology analysis identified disruptions in cell-membrane integrity, flagellar/motility and energy-metabolism pathways, resulting in reduced phenotypic expression, impaired attachment and reduced SRB-induced corrosion (**Supplementary Table S1**). Together, these data support a mechanism in which gTFN couples charge redistribution and nanoscale topology to suppress the gene programs required for stable microbial attachment and biofilm development.

## Microbial biofouling resistance and surface integrity of gTFN

We next assessed whether the charge-engineered gTFN interface improves electrochemical stability under MIC conditions (**see Supplementary Methods S2**). In the presence of OA-G20, gTFN showed greater corrosion resistance than Ni and NiAg. Open-circuit potential (OCP) fluctuations were minimal on gTFN, indicating a more stable electrochemical interface under microbial exposure. In contrast, OCP values for Ni and NiAg fluctuated between −400 mV and −600 mV throughout the exposure period (**Supplementary Fig. S3**), consistent with prolonged microbial activity and evolving corrosion processes.^30^ Nyquist plots further showed that gTFN had higher impedance than Ni and NiAg (**Fig. 2a, Supplementary Fig. S4**). By day 30, gTFN reached the highest low-frequency impedance (|*Z*|□.□□Hz) at 1400 kΩ·cm² ^31^, which was 100-fold higher than Ni (14 kΩ·cm²) and 9-fold higher than NiAg (160 kΩ·cm²). Higher impedance indicates enhanced surface protection and reduced electrochemical charge transfer during corrosion. Electrochemical equivalent circuit (EEC) analysis supported this trend: gTFN showed a charge-transfer resistance (*R*_ct_) of 260 ± 30 kΩ·cm² on day 30, nearly 47-fold higher than Ni (5.6 ± 0.6 kΩ·cm²) and 7-fold higher than NiAg (36 ± 4 kΩ·cm²) (**Supplementary Fig. S3**). Higher *R*_ct_ indicates greater resistance to charge transfer, consistent with reduced anodic dissolution and stronger corrosion protection. Conversely, the lower *R*_ct_ values in Ni and NiAg are consistent with degradation of protective oxide films and increased surface inhomogeneity during prolonged MIC exposure.^32^ Inhibition efficiency (IE) values derived from *R*_ct_^33^ provided further support. NiAg initially exhibited an IE of 67 ± 6%, which declined to 26 ± 3% by day 30 (**Fig. 2b**), whereas gTFN maintained an IE of 98 ± 8% (mean ± s.e.m., n = 3) throughout the exposure period; because IE is bounded by 100%, the upper limit of this interval is truncated at that value. These results indicate that the graphene multilayers and immobilized Ag cap act together to reduce anodic dissolution and restrict microbial access to the Ni substrate.

**Fig. 2.**
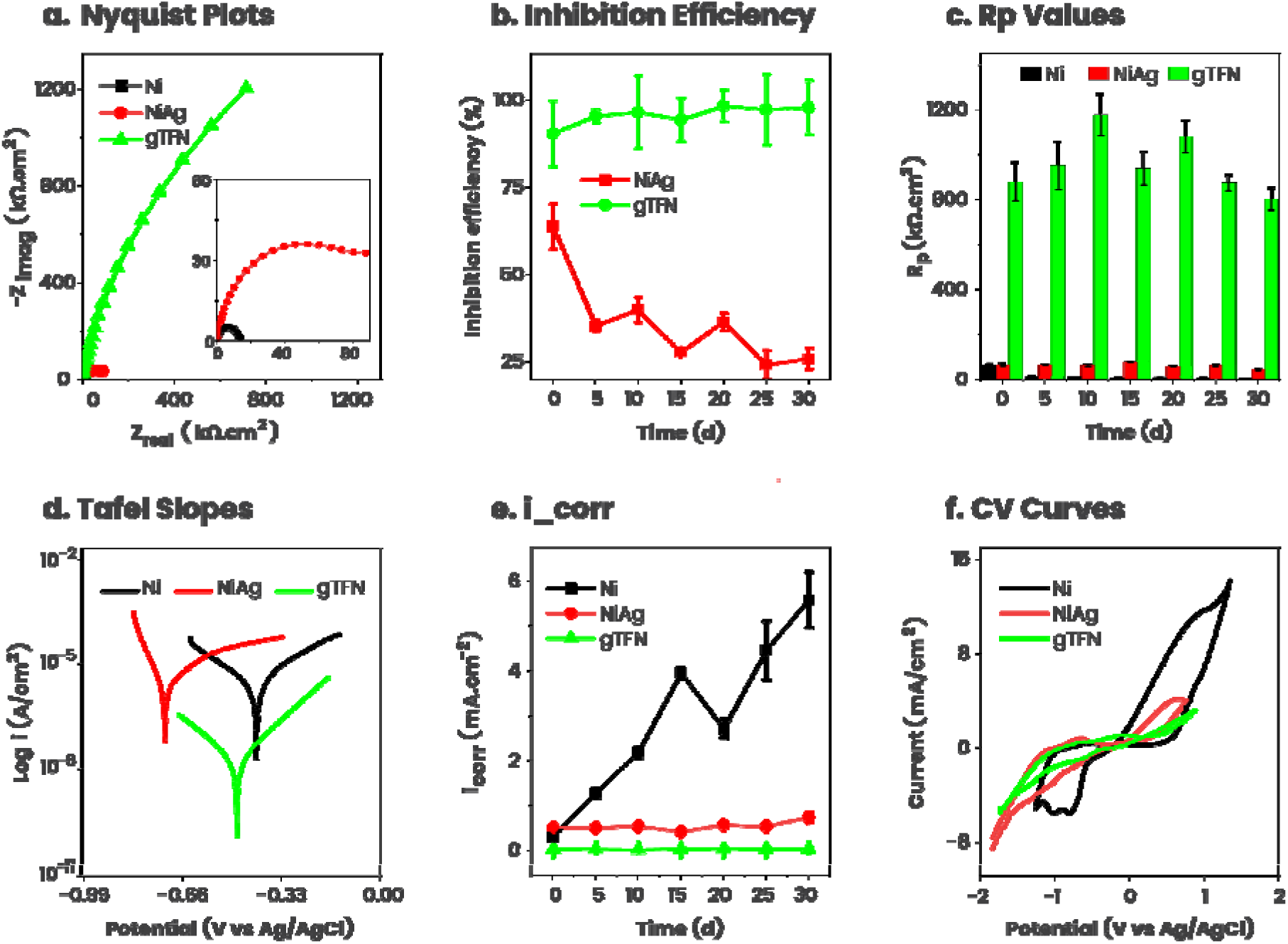
| Electrochemical signatures of charge-governed corrosion resistance in gTFN under SRB attack. Temporal variation of (a) Nyquist plot on 30^th^-day microbial exposure (b) The inhibition efficiency derived from *R*_ct_ values of all samples against nickel (c) Linear polarization derived resistance (*R*_p_) (d) Potentiodynamic polarization plots in a potential range of ±250 mV from the open-circuit voltage (e) Corrosion current (*i*_corr_) (f) Cyclic voltammogram obtained at a scan rate of 0.25 mV s^□^¹ on 30^th^ day. Experiments were performed in triplicate (n□=□3 independent experiments) and analyzed at the 95% confidence level (P□<□0.05). Error bar represent mean□±□s.e.m.

Polarization resistance (*R*_p_) derived from Tafel plots (**Fig. 2c, d**) corroborated the EIS results. On day 30, gTFN exhibited an *R*_p_ of 802 ± 50 kΩ·cm², which is 230-fold and 18-fold higher than Ni (3 ± 0.4 kΩ·cm²) and NiAg (44 ± 5 kΩ·cm²), respectively. The lower *R*_p_ values of Ni and NiAg are consistent with localized microenvironments formed by galvanic coupling between metal sulfides and the underlying metal.^34, 35, 36^ Consequently, gTFN produced a 167-fold decrease in the corrosion current (*i*_corr_) relative to Ni and 22-fold relative to NiAg (**Fig. 2e**). As the Tafel extrapolation on biofilm-covered, non-ideal surfaces carries inherent uncertainty, these i_corr_ values are interpreted alongside the EIS- and LPR-derived parameters, which show the same performance ranking.

Cyclic voltammetry provided additional evidence that gTFN limits microbial redox activity at the surface. Sessile SRB cells on Ni and NiAg can accelerate the corrosion through electron-transfer processes. Cyclic voltammetry analysis (**Fig. 2f**) showed higher cathodic than anodic current densities for both gTFN and NiAg, consistent with SRB-driven biofilm redox activity.^37^ In contrast, Ni exhibited the highest anodic current density (13.3 mA/cm²), which was 13-fold greater than gTFN and indicates greater susceptibility to anodic dissolution. The lower anodic and cathodic current densities on gTFN indicate reduced redox activity and support its resistance to MIC. ^38^

## Phenotypic biofouling and corrosion resistance of gTFN under SRB exposure

We next evaluated whether the electrochemical stability of gTFN translated into reduced microbial attachment, corrosion loss and metabolic activity under SRB exposure. Greater OCP fluctuations are typically associated with electrochemical instability and progressive surface degradation driven by pH shifts, H S generation, metal sulfide precipitation, and variable EPS secretion, all key signatures of MIC. ^39, 40, 41^ Consistent with the minimal OCP variation described above, gTFN showed stable electrochemical behavior under biotic conditions. To evaluate performance under industrially relevant conditions, batch serum bottle reactor experiments were conducted under anoxic conditions (here, “anoxic” denotes an oxygen-free environment, as distinct from “anaerobic” metabolism) using a modified ASTM G31 protocol (**see Supplementary Methods S3 and S4**). After 30 days, corrosion resistance, biofilm formation and elemental dissolution were assessed by weight loss, CFU counts and ICP-MS.

**Figure 3a** summarizes the workflow for DNA extraction, amplification, and sequencing-based analysis. Gravimetric analysis showed that gTFN had the lowest weight loss (8±1 mg/cm²), compared with NiAg (31±4 mg/cm²) and Ni (54±6 mg/cm²). This response correlated with markedly reduced sessile cell densities, which were 29-fold and 11-fold lower on gTFN than on Ni and NiAg, respectively (**Fig. 3b**). The Ag-capped graphene architecture also suppressed biofilm volume (325±30 µm³), achieving 12-fold and 6-fold reductions relative to Ni and NiAg, respectively **(Supplementary Fig. S5)**. Electrolyte analysis indicated limited Ag leaching, with concentrations of 4.6±0.5 µg/L for gTFN and 6.0±0.6 µg/L for NiAg, likely due to Ag S precipitation. Correspondingly, H S levels were also lower in gTFN (3090±400 ppm) than in Ni and NiAg, consistent with reduced SRB metabolic activity (**Fig. 3c**).

**Fig. 3.**
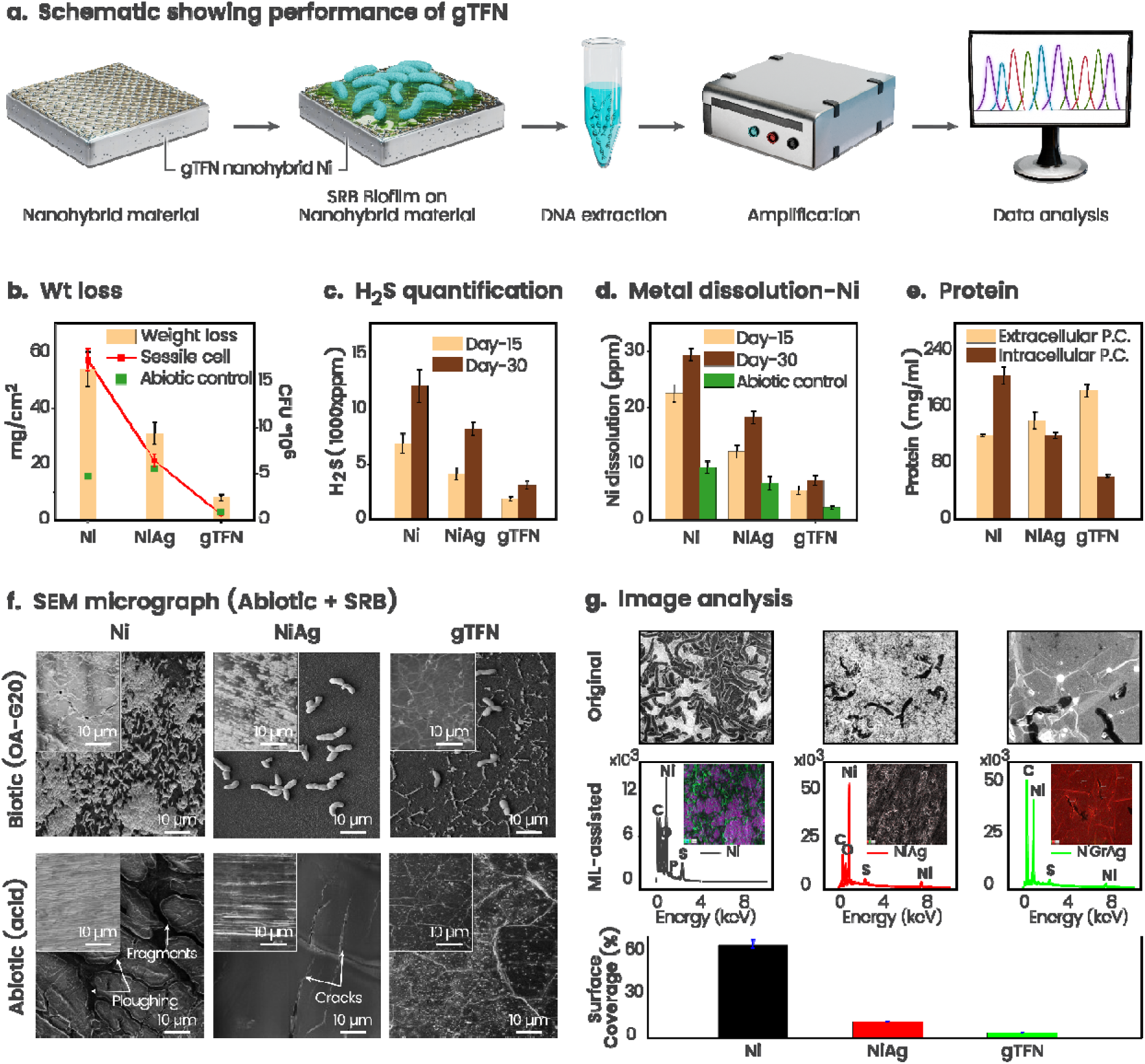
| Microbial biofouling and corrosion resistance of gTFN nanohybrid on Ni: coupled phenotypic and genotypic evidence for charge-governed antifouling. (a) The schematic outlines the experimental workflow from the SRB biofilm over gTFN, followed by DNA extraction, signal amplification, and sequencing-based data analysis. Corrosion resistance performance of different substrates: Ni, NiAg, gTFN, through measuring the correlation between (b) weight loss and sessile cell density over a 30-day interval. (c) SRB metabolic activity through hydrogen sulfide quantification in a serum bottle reactor. (d) MIC resistance through metal dissolution trends. (e) Membrane-disruption tendency and SRB stress response through intracellular (ICP) and extracellular (ECP) protein concentrations. (f) SEM micrographs of different substrates treated in abiotic and biotic (SRB) conditions. Inset micrographs reveal the surface after washing. (g) Machine-learning-assisted image analysis (deep learning) over the SEM micrograph, referred to as “Original”, to measure the surface coverage and affinity of the microbes towards selective sites. All experiments were performed in triplicate (n□=□3 independent experiments) and statistical analyses were carried out at the 95% confidence level (P□<□0.05). Error bars represent mean□±□s.e.m.

Metal dissolution further supported the protective role of gTFN. Ni dissolution from gTFN was 7 ± 1 mg/L, which was 4-fold and 2.6-fold lower than Ni and NiAg, respectively (**Supplementary Methods S5; Fig. 3d**). This reduced dissolution is consistent with limited anodic reactions and lower biocorrosion.^42, 43, 44^ Protein analysis (**Fig. 3e; Supplementary Methods S6**) showed that intracellular protein (ICP) levels in SRB exposed to gTFN were 3.4-fold and 2-fold lower than Ni and NiAg, whereas elevated extracellular protein (ECP) suggested membrane disruption associated with graphene’s serrated topology.^45, 46^ EPS production was also reduced on gTFN (9±1 mg/mL) compared with Ni (29±3 mg/mL) and NiAg (17±2 mg/mL) **(Supplementary Fig. S5)**, supporting inhibition of biofilm maturation.^47, 48^

Topographical analysis further showed that gTFN preserved surface integrity under both abiotic and biotic exposure (see SEM images in **Fig. 3f**). Under abiotic conditions, gTFN remained structurally intact after exposure to 0.5 M H□SO□, whereas Ni and NiAg underwent pronounced acid-induced degradation (**Fig. 3f)**. On Ni, crack formation and microchipping created micropores that can facilitate electrolyte infiltration and accelerate corrosion propagation (**Fig. 3f)**, consistent with higher corrosion current. These observations indicate stronger barrier integrity for gTFN under aggressive chemical conditions.^49^

Under biotic conditions, gTFN also greatly limited SRB colonization. Dense biofilms formed on Ni, whereas microbial attachment on gTFN was sparse. Cells observed on gTFN and NiAg appeared deformed and shrunken (**Supplementary Fig. S6**), consistent with membrane damage associated with Ag thin films and graphene-induced surface interactions.^50, 51, 52^ Machine-learning-assisted image analysis using Cellpose, OTSU and SAM (**Supplementary Fig. S7**) quantified this response, showing only 2.5 ± 0.3% surface coverage on gTFN compared with 67 ± 4% on Ni and 9 ± 1% on NiAg (**Fig. 3g**). Together, these phenotypic, chemical and imaging results show that gTFN reduces microbial attachment, limits Ni dissolution and preserves surface integrity under SRB exposure. ^24, 46, 53, 54^

## Interfacial charge redistribution guides residual microbial attachment

To connect nanoscale charge redistribution with the spatial pattern of residual colonization, we examined partially covered regions where graphene basal planes, wrinkle domains and exposed Ni/Ni□C regions differentially guide microbial attachment; see **supplementary Fig. S1d** for its Raman spectroscopic signatures. Wrinkle domains in gTFN play a role distinct from the antifouling basal plane, acting as localized sites of preferential microbial accumulation, as observed by SEM (**Fig. 4a**). This behavior reflects morphology-driven channeling of SRB colonization rather than uniform surface fouling. Because wrinkle domains occupy only a small fraction of the predominantly basal-plane surface, this channeling concentrates residual attachment at limited sites while the broader graphene-covered regions restrict colonization. This spatial selectivity reconciles localized cell accumulation at wrinkles with the overall 98% reduction in microbial surface coverage.

**Fig. 4.**
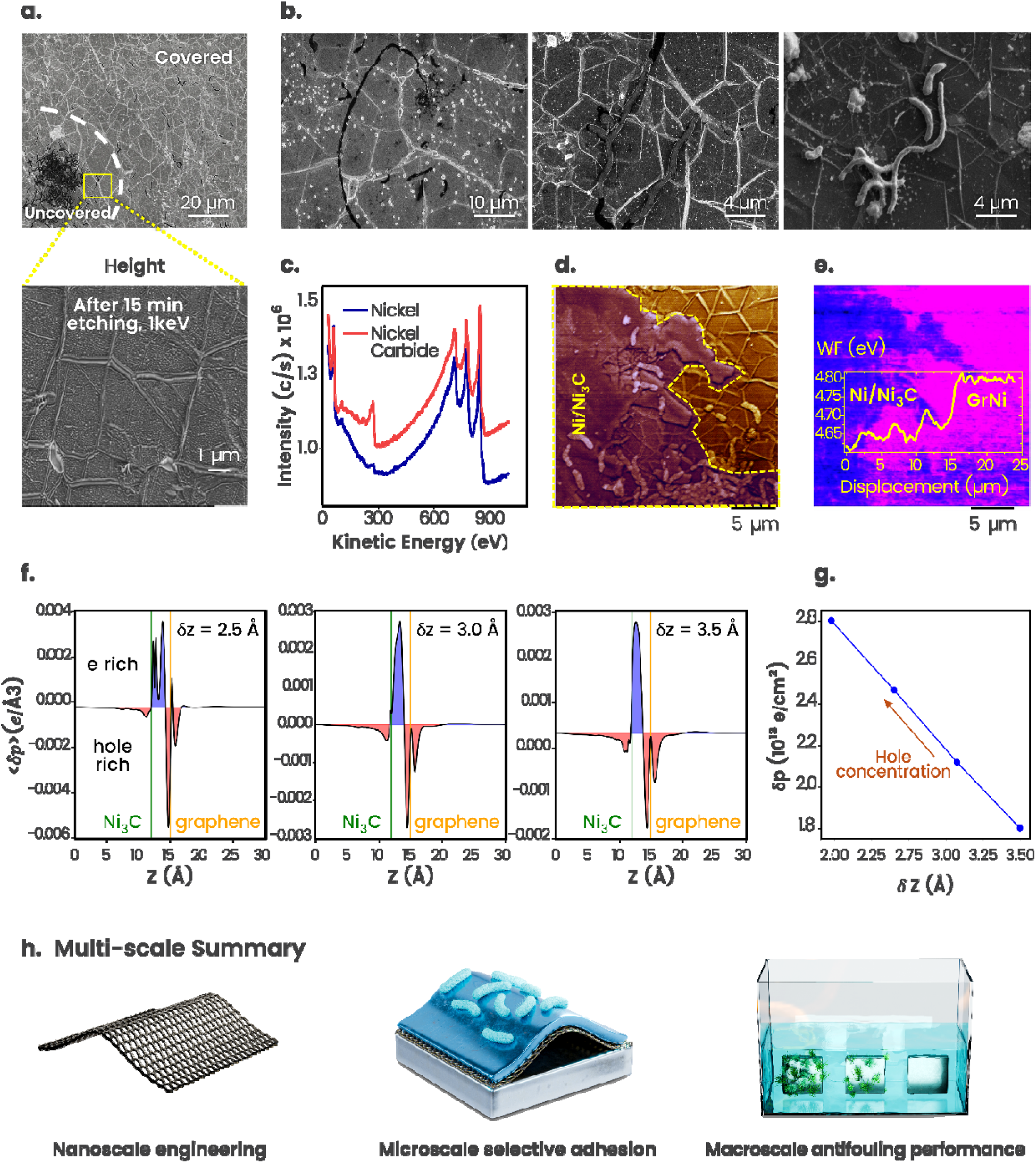
| Interfacial charge distribution and wrinkle-mediated channeling of microbial attachment on gTFN. (a) SEM micrograph of a partially graphene-covered Ni surface. The region inside the dashed line represents etched graphene rich in exposed Ni and Ni_3_C. (b) High-resolution micrographs at different locations reveal tail-to-tail accumulation of microbes over th graphene wrinkles rather than on the basal planes. (c) Auger electron spectra confirm the presence of Ni and its carbide configuration. Concurrently, (d) phase image and (e) surface potential map at fixed lift height at the interface between etched and graphene-covered region reveal work function (eV) distinction between uncoated and covered regions. (f) DFT calculation showing the variation in charge-carrier density (δρ, e Å□³) at separation distances (δz = 2.5 Å, 3 Å and 3.5 Å). Vertical green and yellow lines mark Ni_3_C and graphene. These regions reveal the distribution and density of charge carriers at the interface. (g) Trend showing hole concentration at the separation distance between graphene and Ni_3_C. (h) Schematic summarizing a metric scale depicting nanoscopic solutions of macroscopic performance.

Comparative analysis of etched and graphene-covered regions (**Fig. 4b**) showed that microbial accumulation was greater in etched areas lacking graphene layers, which were primarily composed of Ni/Ni C (**Fig. 4c**). This indicates reduced antifouling protection where graphene coverage was absent. Surface-potential mapping using Kelvin probe force microscopy (KPFM) across the etched and graphene-covered regions (**Fig. 4d, 4e)** revealed a sharp surface-potential contrast. This charge redistribution produced a work function difference (ΔWF = 160 meV; **Fig. 4e inset),** linking electron-rich regions with preferential microbial attachment. Wrinkle geometry with graphene-covered regions generated localized electron-rich sites (**Supplementary Fig. S8**), consistent with earlier observations that defects can influence MIC^3^.

Density functional theory (DFT) calculations support the role of the Ni□C–graphene interface—the covalently bonded contact of the asymmetric metal–graphene–metal (MGM) junction—in shaping this charge landscape (**Supplementary Fig. S9**). Consistent with the KPFM contrast, interfacial charge transfer sets the dipole sign at each contact: electrons accumulate on the Ni side of the buried Ni□C–graphene interface (Niδ□/graphiteδ□), whereas the top graphite–Ag contact, between layers of comparable work function, forms a graphiteδ□/Agδ□dipole. Because the ∼100 nm graphite screens this charge transfer layer-by-layer, the two interface dipoles remain largely decoupled which leads to net charge redistribution in the MGM stacked: Niδ□|graphiteδ□(bottom) … (top) graphiteδ□|Agδ□rather than adding through the film. These calculations establish the interfacial distribution trend and are consistent with the KPFM measurements, which provide the primary experimental evidence. Charge-carrier density (δρ, e Å□³) was evaluated at separation distances of δz = 2.5 Å, 3 Å and 3.5 Å (Fig. 4f). Positive δρ indicates electron accumulation, whereas negative δρ indicates hole accumulation, with the green and yellow lines denoting Ni C and graphene, respectively. At δz = 2.5 Å, a double charge layer forms near graphene (z ≈ 15 Å), with electron density about one-third hole density. As the separation distance increases, hole density decreases (**Fig. 4g**), consistent with the higher electron density observed near rippled or wrinkled graphene in KPFM maps.

In conclusion, SEM micrographs, KPFM imaging, and DFT calculations support a multiscale charge-guided mechanism in which interfacial charge redistribution limits broad microbial attachment, while wrinkle domains locally channel residual cells that reach the surface. This mechanism links nanoscale charge engineering, microscale selective attachment and macroscale antifouling performance under SRB exposure (**Fig. 4h**). gTFN establishes a low-leaching, charge-engineered coating strategy for suppressing microbial attachment and microbially influenced corrosion on Ni. In this architecture, multilayer graphene provides a structurally stable and impermeable conducting scaffold, the immobilized Ag layer as a topping contributes low-leaching antimicrobial activity, and interfacial charge redistribution governs the earliest stages of microbial contact. Electrochemical and phenotypic assays showed a 98% reduction in microbial attachment, a 47-fold increase in charge-transfer resistance and a 167-fold decrease in corrosion current. Transcriptomic profiling indicated suppression of biofilm-associated and energy-metabolism genes, consistent with reduced colonization and metabolic adaptation on gTFN. Together, these findings position gTFN as a charge-engineered antifouling material for controlling microbial biointerfaces in corrosive environments.

## Supplementary Information

Extended protocols, computational parameters and associated references are provided in the “Supplementary Information” of the manuscript as follows:

1. Supplementary Methods (S1 to S13)
2. Supplementary figures (Fig. S1 to Fig. S9)
3. Supplementary Tables (Table S1a to Table S1c)

## Material synthesis and characterization

### Material synthesis

Ni foils of 25 µm thickness and purity 99.99% were purchased from Alfa Aesar (7440-02-0). Multilayer graphene on Ni was prepared in-house using a CVD setup from CVD Equipment Corporation (NY, USA). The quartz tube used is model as First Nano Easy Tube 3000EXT, N1402. After evacuation and Ramp heating (25 to 1000 °C) for 90 minutes, the pressure was around 0.05 Torr. The Ni substrates were subjected to a high-temperature anneal at 1000 ◦C and 500.0 Torr, supported by a carrier gas flow of 349 sccm Ar and 95 sccm H_2_. The active solubility phase was initiated by the introduction of CH4 at a provisional flow rate of 1.98 sccm for exactly 5.53 minutes. During this carbon exposure, the tube temperature was maintained with exceptional stability at 1000.25 ± 0.35 ◦C, while process pressure was held at 500.028 ± 0.357 Torr. To initiate growth and preserve the carbon lattice, the system was subjected to a rapid thermal quench (1000 °C to 650°C). The presence of graphene layers and its distribution are verified from Raman spectroscopy; moreover, commercially purchased Gr/Ni from Graphene Supermarket (CVD-NI-FOIL-2X2-ML) is used to validate the results, especially for wrinkle growth and distribution.

Ni and multilayer graphene nickel foils were ultrasonicated for 10 minutes each in acetone and methanol solutions to remove unwanted contaminants. Ag nanofilms were synthesized on these cleaned foils using pulsed laser deposition (PLD, model: PLD/MBE 2300) equipped with a 248 nm Krypton Fluoride excimer laser. The laser energy/frequency of 350 mJ/10 Hz was used for the deposition. 99.99% pure silver was used as a target and rotated at 30 RPM during the deposition. The substrates (Ni and Gr-Ni foils) were heated to 250 °C and were rotated at 10 RPM. The depositions were conducted at 5 mT pressure, and the substrate-to-target distance was maintained at 96 mm. The depositions were carried out for 3 minutes to maintain a uniform film thickness of ∼3 nm on all the samples, validated through AFM (See Fig. S1a-c).

### Material characterization

Surface characterization was performed using a Helios 5 CX scanning electron microscope (SEM) (Thermo Scientific, Massachusetts, USA) coupled with an Oxford Ultimmax detector (Oxford Instruments, MA, US) and Aztec 5.1 program. Surface roughness was evaluated using VK-X250 laser confocal microscope (Keyence Corp, IL, USA). Raman signatures were analyzed using XploRA PLUS Confocal Raman Microscope (Horiba Scientific, NJ, USA). PHI 710 Scanning Auger electron nanoprobe (Physical Electronics Integrated, MN, USA) was used for additional surface analysis.

### Weight loss and phenotypic response

#### Weight loss, sessile cell count, and nickel dissolution

Weight loss measurements were performed through immersion tests, following a modified version of the ASTM G31 protocol. Sessile cells on the exposed samples were quantified on lactate-C agar plates by serial dilution under anoxic conditions. The concentrations of nickel and silver in electrolytes were determined using an Agilent 7900 Inductively Coupled Plasma Mass Spectrometer (ICP-MS) (Agilent, CA, USA).

#### Protein, exopolysaccharide and H_2_S measurement

The Thermo Scientific™ Coomassie (Bradford) Protein Assay Kit was used to measure intracellular and extracellular protein. Crude EPS was quantified by solvent extraction (**see Supplementary Methods S7**). The concentrations of hydrogen sulfide (H_2_S) in the reactor headspaces were measured using an FD-90A (Forensics Detector, CA, USA), (**see Supplementary Methods S8**.

### Electrochemical corrosion studies

Electrochemical corrosion experiments were performed under two conditions: abiotic and biotic. In both cases, a three-electrode setup was utilized (Gamry Instruments, PA, USA), employing a silver/silver chloride/saturated KCl (Ag/AgCl) reference electrode and a graphite plate as the counter electrode. Abiotic corrosion tests were carried out using 0.5 M H_2_SO_4_ for 24 hours. For microbially influenced corrosion (MIC), pure strains of *Oleidesulfovibrio alaskensis* G20 (10% v/v) were employed, See S3 for details. The growth procedures and lactate-C media used for the corrosion experiments followed the methodology reported in previous studies.^3, 55^

### Biofilm and fouling analysis

#### Microscopy analysis

The morphology of the samples was assessed utilizing the SEM with EDS spectroscopy and analysis of the chemical composition was performed using the Aztec 5.1 program. Biofilm volume on the MIC-exposed samples was measured using confocal laser scanning microscopy (CLSM).

#### Kelvin probe force microscopy (KPFM)

KPFM imaging was performed using a conductive probe (Model: PFQNE_Al from Bruker) and atomic force microscopy (AFM) (Dimension icon scanAsyst). KPFM was carried out in two-pass peak-force mode, where the first pass measures surface topography and the second pass measures surface potential at a fixed lift height. The cantilever was calibrated by thermal tuning, and its work function was measured using a reference Al-Si-Au surface. Under the peak-mode operation, adhesion force (nN) mapping is carried out at 256 x 256 force-distance spectroscopy.^56^

#### Machine learning (ML) analysis

A machine-learning image-segmentation workflow using pretrained Cellpose, Otsu thresholding and the Segment Anything Model (SAM) was used to quantify OA-G20 surface coverage.^57, 58^ ML used SEM images to assess the extent of cell attachment under SRB influence.

#### Pitting profile and corrosion product analysis

Biofilm samples were washed using the ASTM G1-90 protocol before surface-roughness and pitting-profile analysis using CLSM. SEM-EDX and Scanning Auger electron nanoprobe were used for spatial resolution of elemental analysis (**see Supplementary Methods S9** for gradual procedure).

### Transcriptomics analysis

#### Cultivation of OA-G20 under nickel stress and RNA extraction

The pure cultures of OA-G20 were subculture from 1 mL of frozen stock (50% v/v glycerol stock) to 10 mL of freshly prepared modified lactate-C medium^55^. Then the total RNA was extracted using TRIzol Max Bacterial RNA Isolation Kit (Ambion, Life Technologies) by following the manufacturer’s protocol.

#### RNA-seq data analysis

The web-based scientific assessment bioinformatic pipeline called Galaxy was used for data interpretation, as reported in our previous study.^59, 60^. The details of RNA-Seq extraction and data analysis is available at **Supplementary Methods S10 and S11.**

#### Gene ontology, gene clustering and gene co-expression analysis

DESeq2 analysis identified differentially expressed genes, and genes with P < 0.05 and |log□FC| > 0 were selected for further interpretation using the UniProt database (**see Supplementary Methods S12**).

### Density functional theory (DFT)

Calculations were performed in Quantum ESPRESSO using the PBE-GGA exchange– correlation functional, Vanderbilt ultrasoft pseudopotentials and the optB88-vdW dispersion correction, with plane-wave cutoffs of 80 Ry (wavefunctions) and 800 Ry (charge density) and an 11×11×1 Γ-centred Monkhorst–Pack k-point mesh. Non-spin-polarised slabs (≥ 10□) with ≥ 20□ vacuum and a surface-normal dipole correction were relaxed to within 1 meV (energy) and 10 meV□□¹ (force). Plane-averaged charge density and electrostatic potentials at the Ni□C– graphene interface yielded the layer work functions and the interfacial charge-transfer trend (**see Supplementary Methods S13** for full parameters).

## Data Availability

All data supporting the findings of this study are included in the main text and Supplementary Information, which are available online. Any additional datasets or materials generated during the current study will be made available from the corresponding author upon reasonable request.

## Code availability

The image-segmentation workflow used to quantify microbial surface coverage was implemented with the publicly available Cellpose and Segment Anything Model (SAM) packages together with Otsu thresholding; the analysis scripts used in this study are available from the corresponding author upon reasonable request. Density functional theory calculations were performed with the open-source Quantum ESPRESSO package, and all input parameters required to reproduce them are given in **Supplementary Methods S13**.

## Author Contributions

R.D.: conceptualization, investigation, methodology, validation, visualization, writing – original draft. C. S: investigation (Chemical Vapor Deposition), validation (Raman spectroscopy). V.A.S.K.: investigation (pulsed laser deposition), validation. K.J.: investigation (transcriptomics), formal analysis, visualization. B.K.J.: methodology (thin-film deposition), resources, supervision, validation. E.G.: methodology (bioinformatics), resources, software. A.D.: methodology, resources, supervision, writing – review and editing. J.M.: formal analysis (density functional theory), software, visualization, writing – review and editing. P.M.A.: conceptualization, resources, validation, writing – review and editing. M.T.: formal analysis, investigation (Kelvin probe force microscopy), methodology, validation, writing – review and editing. V.G.: conceptualization, funding acquisition, project administration, resources, supervision, validation, writing – review and editing. All authors discussed the results and commented on the manuscript.

## Additional Information

Supplementary information is available in the online version.

## Acknowledgements

The authors acknowledge financial support from the National Science Foundation (OIA-1849206, OIA-1920954 and 2418752), National Institutes of Health (#5P20GM103443-20), and Civil and Environmental Engineering Department at South Dakota Mines. We thank Dr. Suvarna Talluri for technical guidance on serum bottle experiments. All the authors thank Dr. Takshashila Tripathi for the in-depth discussion on transcriptomics analysis.

**Correspondence** and requests for materials should be addressed to Venkataramana Gadhamshetty.

